# Metabolic profiling and genome-scale modelling uncover mechanistic drivers of microbiome stability in synthetic maize root community

**DOI:** 10.64898/2026.05.20.726550

**Authors:** Jenna Krumbach, Lisa Schönherr, Patrizia Kroll, Vera Wewer, Sabine Metzger, Till Ischebeck, Martina Feierabend, Nadine Töpfer, Stanislav Kopriva, Richard P. Jacoby

## Abstract

Stability is a desirable property for agricultural microbiomes, but there is a poor understanding of the mechanisms that mediate microbial community stability. A representative bacterial synthetic community from maize roots has been proposed by Niu et al. (2017, PNAS, 114:E2450) as a model system to study microbiome stability. This SynCom assembles stably when all seven members are present, but community diversity collapses without the keystone *E. ludwigii* strain. In this study, we used complementary *in vitro* experiments and *in silico* metabolic modelling to assess the role of metabolites for the stability of this SynCom, by defining the metabolic niches occupied by each strain, as well as their cross-feeding phenotypes and B-vitamin dependencies. We show that the individual member strains occupy complementary metabolic niches, measured by the depletion of distinct metabolites in exometabolomic experiments, as well as contrasting growth phenotypes on diverse carbon substrates, patterns which are largely recapitulated by computational simulations. Minimal medium experiments show that the established seven-member community comprises a mixture of prototrophic and auxotrophic strains. Correspondingly, experimental and *in silico* cross-feeding phenotypes showed that spent media harvested from the prototrophic strains can sustain growth of two auxotrophs and let to the identification of B-vitamin dependencies. Altogether, this study highlights the complementary power of *in vitro* and *in silico* approaches and suggests that the metabolic mechanisms of this SynCom can serve as design principles to inform the rational assembly of stable plant-associated microbial communities.

## Introduction

A deeper understanding of community stability is a major research goal in the field of microbiome science and technology [1]. Potentially, stability criteria could help to predict how different microbiomes respond to disruption, such as environmental change or biotic invasion [2]. Furthermore, a mechanistic understanding of microbiome stability can help to rationally formulate bio-inoculants in agriculture and medicine, via combining cooperative strains into stable synthetic communities that will persist in the target environment [3].

Conceptually, metabolism is positioned as a central factor mediating microbiome stability, underpinned by two major processes: 1) niche differentiation, and 2) cross-feeding [4,5]. The ecological principle of niche differentiation stipulates that organisms which coexist in the same habitat avoid competition by consuming different resources [6]. Although theoretical studies predict that niche differentiation plays a major role in facilitating the diversity of microbiome composition, there are still significant knowledge gaps regarding the specific metabolites consumed by individual strains [7]. Cross-feeding facilitates microbial diversity, because the metabolites released by one strain can support the coexistence of other strains, e.g. by supplying vitamins and amino acids that nourish auxotrophs [8]. However, there is a relatively poor mechanistic knowledge of which microbial strains engage in cross-feeding, and what specific metabolites are shared [9]. The laboratory study of metabolic interactions in microbial communities has been hampered by the lack of experimentally tractable systems [10]. However, a previously characterised seven-member bacterial synthetic community (SynCom) from maize roots is emerging as a model system for plant microbiome research [11]. This community comprises Proteobacteria, Bacteroidetes and Actinobacteria phyla: *Stenotrophomonas maltophilia* (Sma), *Brucella pituitosa* (Bpi), *Curtobacterium pusillum* (Cpu), *Enterobacter ludwigii* (Elu), *Herbaspirillum robinia* (Hro), *Chryseobacterium indologenes* (Cin) and *Pseudomonas putida* (Ppu). Notably, this SynCom assembles stably when all seven members are present, but the removal of strain Elu results in a collapse of microbial community diversity [11]. This observation is reminiscent of the keystone effect in macroecology, whereby one species plays a disproportionately large role in facilitating ecosystem diversity [12].

In recent years, computational models that capture the metabolic interactions between the host and its associated microbiota have emerged as a powerful tool to understanding these complex multi-organismal metabolic interactions. Genome-scale metabolic models of individual microbes can be reconstructed from the genome sequence and encompass the metabolic networks of each organism. Simulations of metabolic fluxes of individual members on different growth media as well as within the community can provide mechanistic understanding of resource use patterns and metabolic exchanges between interacting community members [13].

In this study, we describe the metabolic characteristics of this previously established seven-member bacterial SynCom through *in vitro* and *in silico* experiments. To this end, we 1) Assess the metabolic niche of each strain, 2) Describe cross-feeding interactions between donor and recipient strains, and 3) Define the B-vitamin responses of SynCom members (see Fig.1 for an overview of our combined *in vitro* and *in silico* approach).

**Figure 1:**
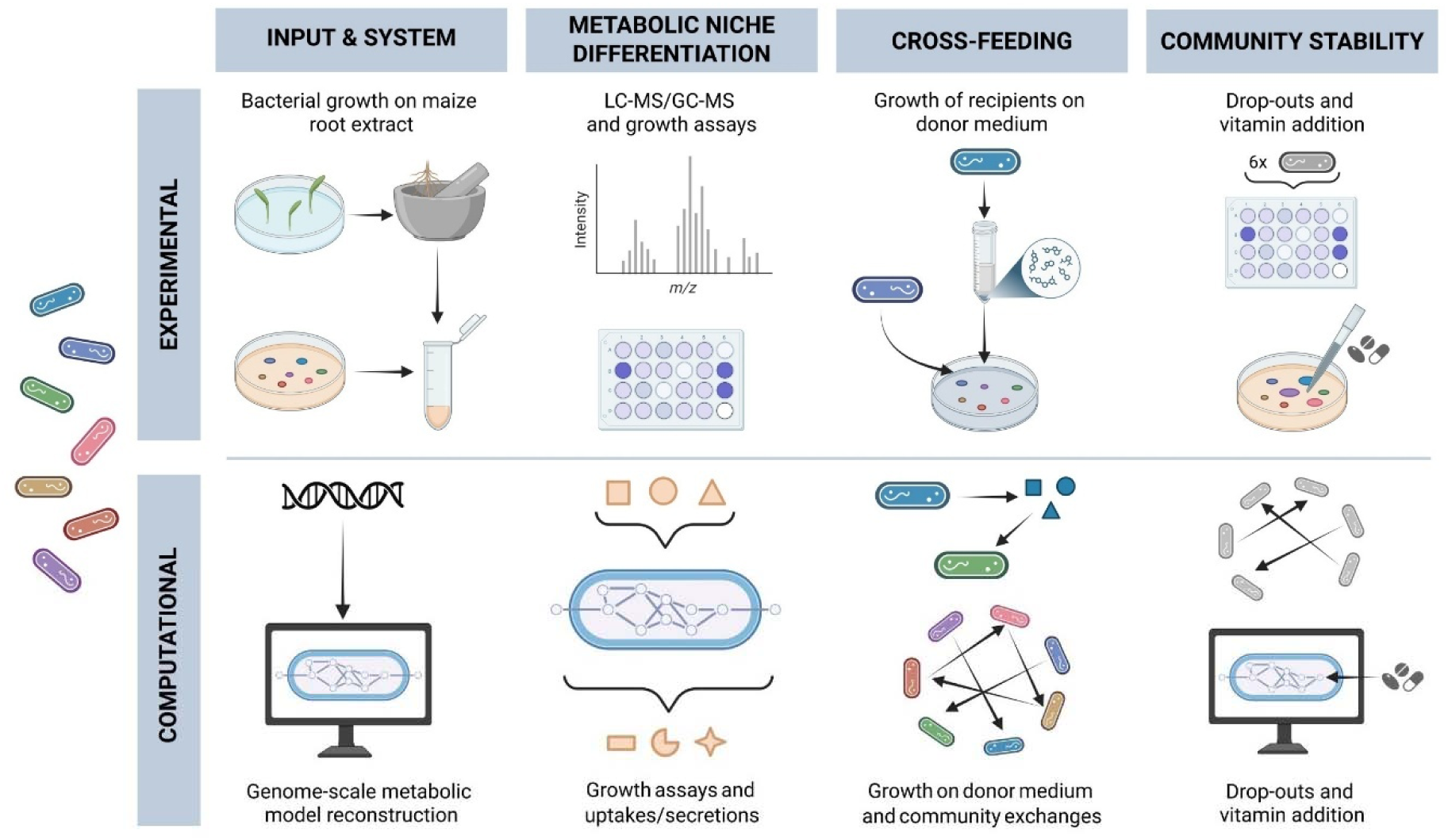
Overview of the combined experimental and computational framework used to identify metabolic characteristics of a seven-member bacterial SynCom. In this study we characterise the metabolic properties of a previously established seven-member bacterial SynCom using complementary experimental and computational approaches. We assessed the metabolic niche of each strain, identified cross-feeding interactions between donor and recipient strains, and determined community responses to B-vitamin availability. Genome-scale metabolic modelling was used alongside *in vitro* experiments to simulate growth, metabolite exchange, and community perturbations under defined conditions. Together, these analyses provide mechanistic insight into how metabolic traits and dependencies shape community assembly and stability. Figure created with BioRender.com.

## Methods

### Exometabolomics of bacteria cultivated on maize root extract

Microbial exometabolomics on maize root extracts was conducted according to the method adapted from Jacoby et al [14]. Maize seeds (v. Sunrise) were purchased from Agri-Saaten Ltd, and sterilised in 70% ethanol for 15 min, followed by 5% NaHClO_4_ for 15 min, rinsed five times with sterile water, then incubated in sterile water at 50° C for 10 min. Individual seeds were then placed in Petri dishes with 10 mL of sterile water and incubated in the dark for 4 days at 30° C. Next, germinated seedlings were transferred into a sterile hydroponic growth system, enclosed in transparent plastic boxes with HEPA-filters for gas exchange (Sac O_2_ Ltd), with roots supported by 3 mm glass beads, and growth medium containing 0.5× Hoagland’s solution. Plants were placed in a growth chamber with 23/18° C day/night temperature, 16 h day length, and 150 µmolm^-1^s^-1^ light intensity. Plants were cultivated for seven days, with growth medium changed once. At harvest, roots were separated from shoots, washed three times in sterile 10 mM MgCl_2_, then root tissue was snap-frozen in liquid N_2_.

For metabolite extraction, frozen maize root tissue was ground to a fine powder using a mortar and pestle and liquid nitrogen. Next, 200 mg of tissue powder was placed into a 1.5 mL tube and incubated with 1 mL of 90% MeOH at 60° C for 30 min with 1500 rpm shaking. Tubes were centrifuged at 10,000× g for 10 min, then 800 µl of supernatant was transferred to a new tube and dried down in a vacuum centrifuge overnight. Next, dried metabolites were dissolved in water and filter-sterilised (0.22 µm pore size). Total organic carbon (TOC) concentration was measured using a Dimatoc 2000 (Dimatec).

For bacterial pre-culture, strains were streaked from glycerol stocks onto TSA plates (0.5× TSB, 1.2% agar) and incubated at 28° C for 1-2 days. Individual colonies were picked and transferred into 4 mL of 0.5× TSB at 28° C with 200 rpm shaking for 1-2 days. Cells were harvested by centrifuging 900 mL of culture at 5,000× g for 5 min at RT. Cell pellets were then washed twice with 900 mL of sterile 10 mM MgCl_2_ and resuspended at a final OD600 of 1 in sterile 10 mM MgCl_2_. The C7 mixture of all seven strains was prepared by equally combining washed bacterial cells at OD=1.

Next, bacteria were cultivated on an M9 growth medium where maize root extracts were the sole carbon source. The medium contained 24 mM Na_2_HPO_4_, 20 mM NH_4_Cl, 11 mM KH_2_PO_4_, 4 mM NaCl, 1 mM MgSO_4_, 100 μM CaCl_2_, 50 μM Fe-EDTA, 50 μM H_3_BO_3_, 10 μM MnCl_2_, 1.75 μM ZnCl_2_, 1 uM KI, 800 nM Na_2_MoO_4_, 500 nM CuCl_2_, and 100 nM CoCl. To this, maize root extracts were added to a final concentration of 720 μg C per mL. Growth assays were conducted in a 48-well plate, by adding 20 µL of resuspended bacterial pellet into 380 µL of medium, and incubating the plate in a plate reader (Tecan Infinite Pro 100) at 28° C for 24 h, with shaking (3 min continuous orbital shaking followed by 7 min stationary, shaking amplitude 3 mm). A negative control (no bacteria) was also prepared by adding 20 uL of sterile 10 mM MgCl_2_ to the growth medium and was incubated side-by-side with the bacterial cultivations. Spent media were harvested by centrifuging cultures at 10,000x g for 3 min, then filter-sterilising the derived culture supernatants using 0.22 µm spin filters.

For untargeted LC-MS analysis of culture supernatants, 5 μl of sample was loaded onto a C18 column (XSelect HSS T3, 2.5 µm particle size, 100 Å pore size, 150 mm length, 3.0 mm inner diameter; Waters), using an HPLC (Dionex Ultimate 3000, Thermo Scientific). Solvent A was 0.1% formic acid (FA) in water, solvent B was 0.1% FA in methanol, and flow rate was 500 μl/min. Samples were eluted using the following gradient: hold at 1% B between 0 to 1 min, linear increase to 40% B until 11 min, linear increase to 99% B until 15 min, hold at 99% B until 16 min, linear decrease to 1% B until 17 min, and finally, hold at 1% B until 20 min. MS analysis was conducted using a Q-TOF MS (maXis 4G; Bruker Daltonics), following electrospray ionization (ESI). The MS was operated in both positive and negative-ion mode, using N_2_ as drying gas at a flow rate of 8 litres/min, dry heater set to 220° C, nebulizer pressure of 1.8 bar, capillary voltage of 4,500 V, and collision radio frequency voltage of 370 V (mass range of 50 to 1,300 m/z). Scan rate was 1 Hz.

To process the untargeted LC-MS data, raw.D files were centroided and converted to .MZMXL using the MSConvert program [15]. Files were then uploaded to XCMS online [16] and were aligned using the following parameters: m/z tolerance of 15 ppm, minimum peak width of 10 s, maximum peak width of 60 s, signal/noise threshold of 10, overlapping peaks split when m/z difference was greater than 0.01 m/z, features only considered if they occurred in at least five consecutive scans with an intensity greater than 5,000. Also, features were only considered if they occurred in at least two of three replicates from any sample group. The CAMERA algorithm was implemented to detect isotopes and adducts. Following export of files from XCMS online, data from both positive and negative-ion modes were combined, and filtered to remove isotopic peaks and filtered to only include MS features with RT between 1 and 16 min.

The statistical analysis of the LC-MS exometabolomic data had two aims: 1) To detect maize metabolite ions that were depleted from the medium following bacterial growth, and 2) To detect microbe-derived metabolite ions that were enriched in the medium following bacterial growth. Both strategies involved comparing the metabolomic profiles of the inoculated samples versus the uninoculated negative control. For a metabolite ion to be categorised as depleted, thresholds were signal intensity < 50% versus sterile control, p-value < 0.05. For a metabolite ion to be categorised as enriched, thresholds were signal intensity > 200% versus sterile control, p-value < 0.05.

For GC-MS quantification of bacterial primary metabolite depletion from maize root extracts, 50 µl of spent or fresh medium was pipetted into 500 µl of methanol/chloroform/water (5:2:1 (v/v/v)), and 200 of *allo*-inositol (5 µg/ml) was introduced into each sample as an internal standard. Next, 100 µL of the polar fraction was dried under a stream of N_2_ gas and derivatised with 15 µl methoxyamine hydrochloride in pyridine (30 mg/ml) and 30 µl N-Methyl-N-(trimethylsilyl) trifluoroacetamide (MSTFA) [17]. The samples were analysed on an Agilent 5977N mass selective detector connected to an Agilent 7890B gas chromatograph, as previously described [18]. Primary metabolites were quantified according to the intensity of reporter ions previously obtained for pure reference compounds, normalised to the intensity of the *allo*-inositol internal standard. Bacterial depletion of primary metabolites was measured by comparing the normalised metabolite abundance in the inoculated samples versus the sterile controls.

### Phenotype microarray

Phenotype microarrays were conducted using BIOLOG EcoPlate according to the manufacturer’s instructions. The seven bacterial strains were pre-cultured, harvested and washed as described above, then diluted to an OD value of 0.1 in sterile 10 mM MgCl_2_. At this point, strains were mixed equally mixed together to compose either C7 communities or C6 drop-outs, and 150 µL of bacterial suspension was inoculated into each well of the 96 well-plate. Plates were incubated at 28° C for two days, and A590 was measured using a Tecan M100 Infinite Pro.

For analysis of phenotype microarray data, A590 values from all substrate-containing wells were compared to a negative control (no substrate) via Student’s t-test, and growth was considered positive if p-value < 0.05 and A590 > 0.1. In our hands, two substrates (2-hydroxy-benzoic acid and alpha-D-lactose) were not utilised by any bacterial inoculation and were therefore excluded from the analysis.

### Minimal medium growth assays

For assays of bacterial growth on single carbon substrates, bacteria were pre-cultured, harvested and washed as described above. Medium composition was the same as the M9 minimal medium described above, where all individual carbon sources were included at a concentration of 720 µg C per mL. Growth assays were conducted in a 96-well plate, where each well contained 95 µL of medium, inoculated with 5 µL of bacterial suspension, and cultivated in a plate reader (Tecan M100 Infinite Pro) using the same program as described above, except that data was collected over two days. Growth curves were quantitatively analysed using the Growthcurver R package [19].

### Bacterial cross-feeding assays

To analyse pairwise cross-feeding phenotypes across different carbon sources, the assay began by pre-culturing the three donor strains which could successfully grow on minimal medium (Elu, Hro and Ppu). This was conducted by picking colonies from TSA plates and transferring them into 4 mL of minimal medium, formulated as described above with either glucose, malate or alanine as sole carbon source at 720 µg C per mL, then incubating at 28° C with 200 rpm shaking for 2 days. The derived culture supernatants were harvested by centrifuging at 10,000x g for 3 min, then filter-sterilising the supernatants using 0.22 µm filters. In parallel, all seven recipient strains were pre-cultured on 0.5×TSB, harvested and washed as described above. In a 96-well plate, 5 µL of bacterial suspensions (OD=1) from the recipient strains were inoculated into 95 µL culture supernatants harvested from the donor strains, alongside a negative control of sterile 10 mM MgCl_2_ that was included to check for sterility of culture supernatants. Growth assays and quantitative analysis were performed as described above.

### Vitamin response assays

Several methodological strategies were used to dissect the vitamin responses of these strains. First, the responses of all seven strains to a mixture of B-vitamins were undertaken, whereby bacteria were pre-cultured, harvested and washed as described previously, and inoculated into a minimal medium containing either all eight B-vitamins or no vitamin addition. The provided vitamins were: thiamine, riboflavin, nicotinamide, pantothenate, pyridoxine, biotin, folate, cobalamin. In this first experiment, the carbon sources were a mixture of 20 carbon substrates (glucose, myo-inositol, sorbitol, sucrose, xylose, 2-oxoglutarate, citrate, malate, pyruvate, succinate, GABA, glutamate, glycine, L-alanine, D-alanine, β-alanine, L-arginine, D-arginine, putrescine and urea). Each carbon source was provided at 36 mg C per L, such that the combined carbon concentration in the medium was again 720 mg C per L. These diverse carbon substrates were provided to maximise the probability of observing a growth phenotype, unobscured by substrate preference effects. Next, we investigated the specific vitamins required for the growth of strains Bpi and Cpu, this time providing single carbon sources of either alanine (for Bpi) or glucose (for Cpu). These substrates were chosen following preliminary experiments that identified these compounds as preferred single carbon substrates for each strain. Here, two approaches were used to pinpoint vitamin auxotrophies: either the addition of a single vitamin (V1 approach) or the removal of a single vitamin from the eight-vitamin mix (V7 approach). In both approaches, vitamins were provided at the previously used concentrations.

### Metabolic network reconstruction and flux modelling

Genome-scale metabolic models for all seven SynCom members were reconstructed from published genomes [20] using CarveMe 1.6.2 [21]. Models were curated following standard practices [22] with particular focus to mass and charge balancing, removal of duplicate reactions, and gap-filling of growth-associated pathways (see SI for details of the curation process). Model quality was evaluated using MEMOTE 0.17.0 [23], yielding an average overall quality score of 86.4% ± 0.5 for all seven models. In each model, growth maximisation was set as the objective using the biomass objective function. All simulations were performed in Python using COBRApy 0.29.1 [24]. Flux solutions were obtained using parsimonious flux balance analysis (pFBA) [25] or flux variability analysis (FVA), [26, 27] under defined medium conditions. Unless stated otherwise, simulations were conducted in a medium composed of primary metabolites identified in maize root extract by GC–MS (Table S1), supplemented with all compounds present in the M9 medium described above (termed “*in silico* maize root medium” Table S2). When required, alternative media reflecting specific *in vitro* experimental conditions were applied as described in the following paragraph.

Exometabolomic simulations were conducted using pFBA to analyse uptake and secretion profiles of all models grown on *in silico* maize root medium. Phenotype microarray simulations were conducted for carbon sources from BIOLOG EcoPlates that were present in the models. Growth was evaluated through pFBA in a simulated M9 medium containing a single carbon source. Similarly, minimal-medium growth assays were simulated using M9 medium combined with one of nine carbon sources to identify donor species. For cross-feeding analyses, secretion profiles of donor species simulated on this minimal medium were determined using FVA. To ensure that these profiles represent a near-optimal growth state of the donor, COBRApy’s flux_variability_analysis() function was run with the parameters fraction_of_optimum = 0.98 and pfba_factor = 1.05, requiring the objective value to reach at least 98% of its maximum while allowing the total flux to exceed the pFBA solution by a maximum of 5%. The resulting profiles were then used as growth media in pFBA simulations for all seven strains. Responses to B-vitamins were assessed with pFBA by supplementing vitamins to a M9 medium containing 20 distinct carbon sources (see vitamin response assays section). Additional simulations examined V1 and V7 conditions for Bpi using alanine as the sole carbon source.

Models for the C7 community and C6 drop-outs were constructed using the Community() function in MICOM 0.37.1 [28] with individual strain models provided as input without predefined abundances. The community objective was defined as maximisation of the summed growth rates of all members. The trade-off fraction was set to 1. This parameter ranges from 0 to 1 and limits community growth to a chosen fraction of its maximum, thereby influencing how growth is distributed among individual members. All community analyses were performed using pFBA, implemented via MICOM’s cooperative_tradeoff() function. The C7 community was used to simulate the exometabolomic and BIOLOG EcoPlate experiments described above, and BIOLOG EcoPlate simulations were additionally repeated using the C6 drop-out models. Cross-feeding interactions within the C7 community were further analysed by examining metabolite exchange between members using pFBA. The code needed to reproduce the results of this paper can be found at https://github.com/Toepfer-Lab/C7-maize-syncom.

## Results

### Exometabolomic profiling of the SynCom on maize root extract

First, we hypothesised that the stable assembly of this community could be the result of metabolic resource partitioning, whereby the strains avoid direct competition by occupying differential metabolic niches in the maize rhizosphere habitat. To investigate this, we undertook a high-throughput LC-MS exometabolomic analysis to identify which maize root metabolites the strains consume as growth substrates. Methodologically, this involved cultivating the microbes on a growth medium where maize root extracts were the sole carbon source, then analysing the derived culture supernatants using LC-MS. The derived data enabled us to profile how the strains had modulated their chemical environment, providing new information about the maize root metabolites consumed by each strain, and the microbe-derived metabolites released into the growth medium.

Our data analysis strategy initially focussed on characterising microbial substrate preferences, by identifying the metabolite peaks that were present in the sterile medium but depleted following bacterial growth. The derived data is presented as a heatmap of 425 metabolite ions depleted by at least one strain (Fig 2a, Table S3). Investigating the metabolite depletion profiles, we observed clear evidence of niche differentiation, because each strain depleted a distinct set of metabolite ions. Intriguingly, the C7 seems to combine the metabolic capabilities of each individual strain, and due to this ‘addition effect’ the C7 occupies a much broader metabolic niche compared to any single individual.

**Figure 2:**
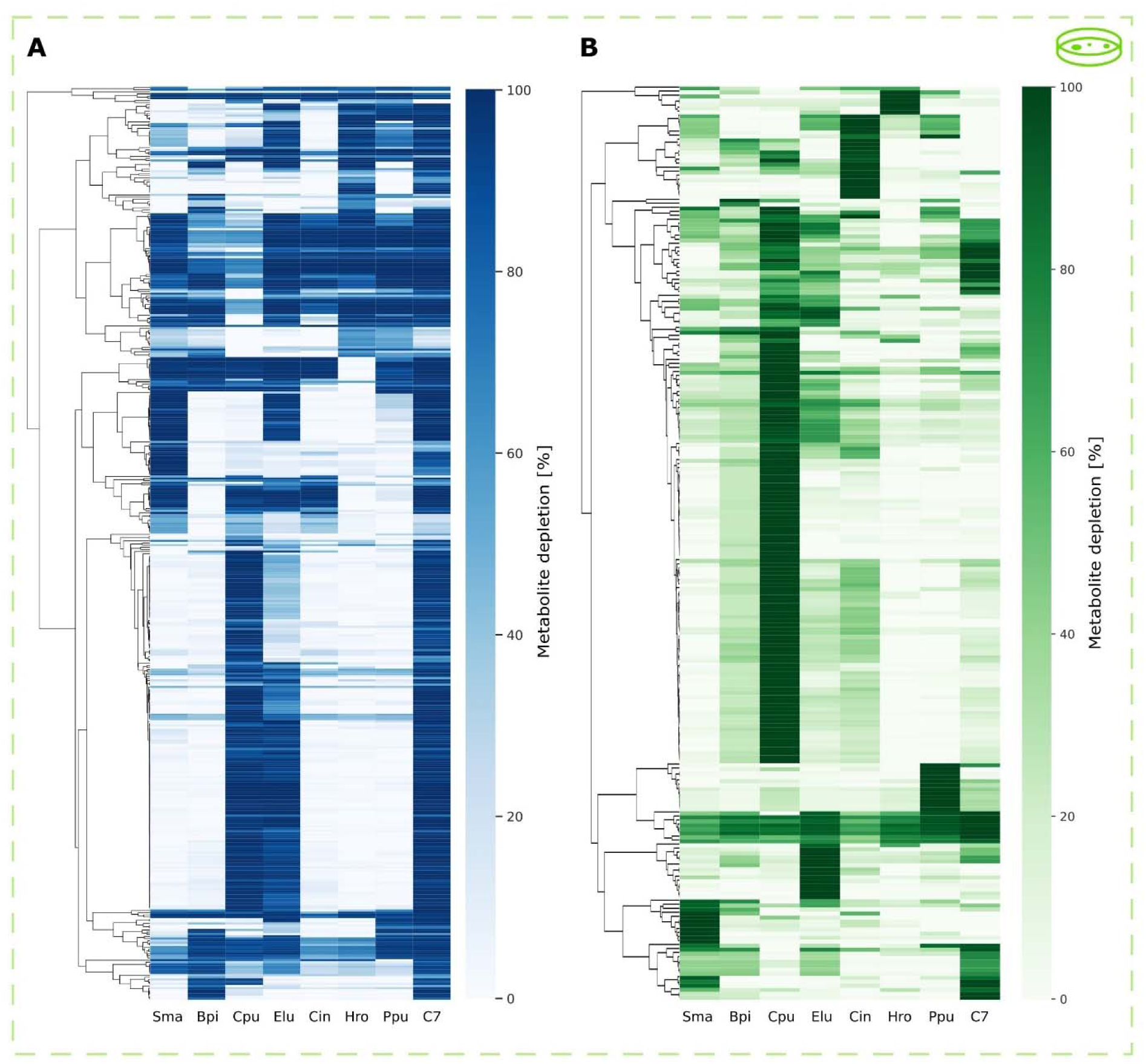
Metabolic footprints of the SynCom and constituent strains on maize root extract. (A) Heat map showing metabolite depletion profiles for 425 metabolite ions present in maize root extract, measured by LC-MS exometabolomics. Metabolite ions were included if they were depleted by at least one inoculum (LC-MS abundance <50% compared to sterile control, p<0.05), cell colour represents the mean depletion value from three independent replicates. Rows were clustered via Pearson correlation. (B) Heat map of metabolite enrichment profiles for 228 metabolite ions that exhibited higher abundance in the inoculated samples versus sterile controls (abundance >200%, p<0.05). Cell colour represents the mean enrichment value from three independent experiments, measured via untargeted LC-MS exometabolomics. Rows were clustered via Pearson correlation.

The heatmap is dominated by a large grouping of depleted metabolite ions that were only depleted by three inocula: Elu, Cpu and C7. It has to be noted that our aim was not to generate a comprehensive list of all detected metabolites, but instead to narrow down which metabolites niches are occupied by the individual strains. To this end, we sought to gain more information about the chemical identity of these depleted metabolite ions (see Supplementary Information). To present the data, we integrated the heatmap of metabolite depletion profile with the chemical categorisations of the candidate metabolite IDs. Chemically, many of these m/z values were categorised in the group ‘glycosylated compounds’, and closer inspection of the data indicated that this cluster includes a large number of maize secondary metabolites, including sugar conjugates of flavonoids and benzoxaolones (Table S3). The overlapping niches of these two strains is notable, because the work of Niu et al (2017) found that the absence of Elu allows Cpu to dominate the assembled community. The heatmap also contains a relatively small cluster of 52 metabolite ions that were universally depleted by all seven strains. Chemical categorisation showed that this cluster contained a large proportion of primary metabolites, such as amino acids and organic acids.

Our second data analysis strategy with the untargeted LC-MS exometabolomics dataset involved determining which microbe-derived metabolites were released into the growth medium by each strain. This heatmap is dominated by a large cluster of metabolite ions predominately released by strain Cpu (Fig 2b). Chemically, many of these m/z values were categorised as ‘amino acids and derivatives’ and include several free benzoxazolones lacking the sugar group (Table S4, Figure S1a). Due to Cpu’s potential to dominate the other strains, we suggest that these metabolite ions could represent antimicrobial compounds or other chemical antagonists. We extended this analysis by GC-MS exometabolomic profiling of the same samples in which 25 primary metabolites were identified (Table S1). This analysis showed that the consumption of major sugars, organic acids and amino acids was common for all seven strains (Fig 3a).

**Figure 3:**
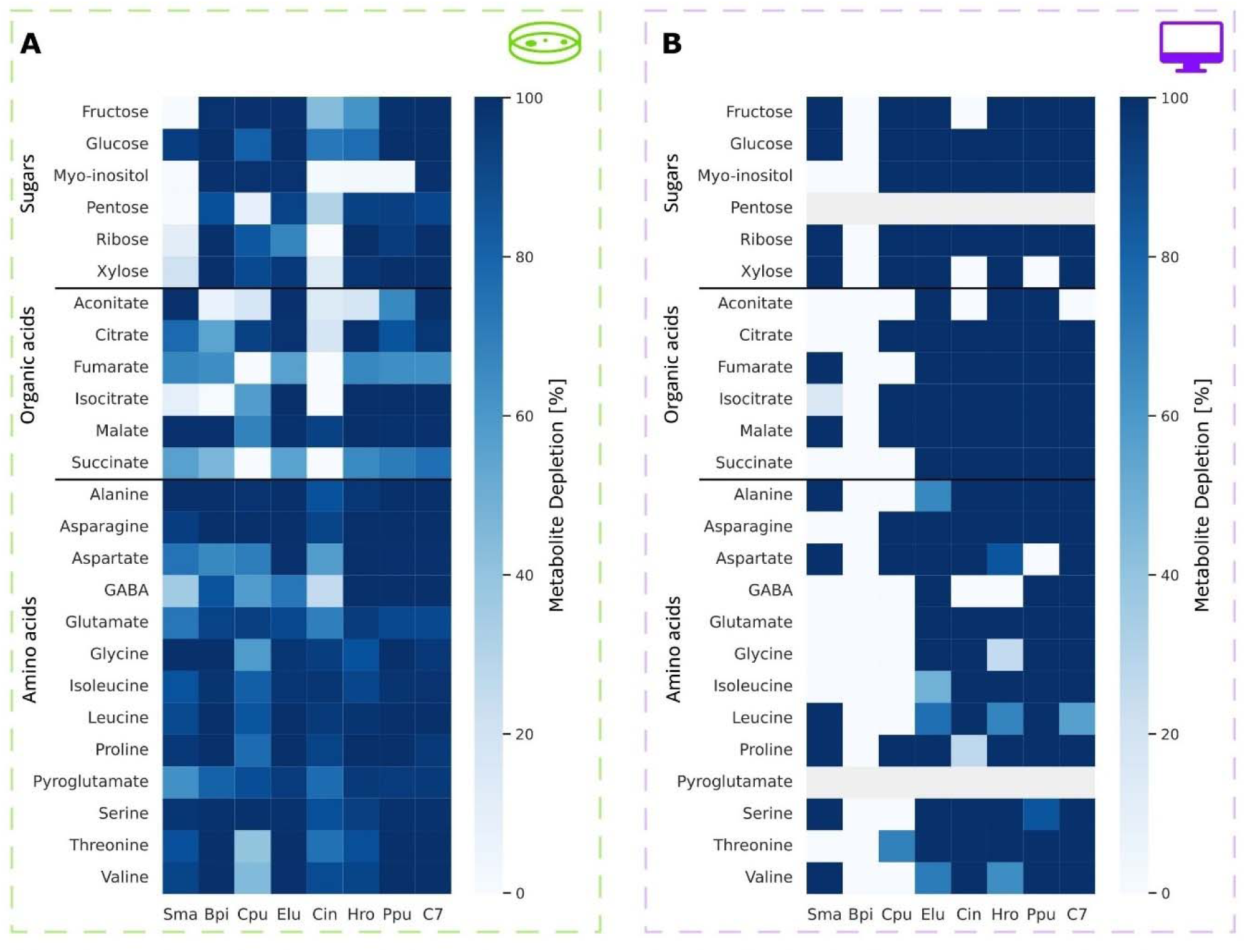
Depletion of primary metabolites from maize root extract by the SynCom and constituent strains. (A) Heat map of primary metabolites depleted by the SynCom strains measured via GC-MS exometabolomics of SynCom strains cultivated on maize root extract. Cell colour represents mean metabolite depletion value of three independent replicates. (B) Heat map of primary metabolites depleted by the SynCom strains simulated on a medium consisting of those primary metabolites together with a M9 medium. Cell colour represents the metabolite depletion predicted with pFBA, calculated as the absolute flux of the corresponding exchange reaction normalised by its upper bound. A grey cell colour represents no predicted growth as pentose and pyroglutamate were not available in the models. The mean growth rate of all seven bacteria in the community is used for the C7 scenario.

In a complementary approach, we generated genome scale metabolic models for the 7 strains and the C7 community and used them for a flux balance analysis. For the simulations, the 25 primary metabolites together with the M9 medium were used as *in silico* maize root medium (Table S5). It shall be noted that none of the metabolic models includes exchange reactions for pentose and pyroglutamate and thus these metabolites could not be used by the models. Our simulations largely recapitulated the experimentally observed depletion patterns, with the majority of these metabolites being consumed by most strains (Fig 3b). Together, this indicates that primary metabolites constitute broadly shared resources rather than strain-specific niche factors and that primary metabolites likely play a minor role in mediating niche differentiation.

Only for Bpi we observed a notable discrepancy between the *in vitro* and *in silico* results, since the model predicted neither growth nor metabolite uptake on the defined medium. This observation could be due to the lack of essential metabolites in our *in silico* maize root medium which are present in the maize root extract.

Based on the limited number of metabolites in the *in silico* maize root medium, the number of predicted released compounds was noticeably less than *in vitro*, 13 in total (Supplementary Fig S1b). Among all individual strains, Cpu released the most metabolites. Half of them were unique to this strain, reflecting the pattern observed in the LC-MS-based enrichment profile. Notably, one of these secreted metabolites was 3-hydroxydecanoic acid which is known for its involvement in plant immune response and antifungal properties [29].

### Substrate utilisation profiles of the SynCom and C6 drop-outs

The identification of specific chemical structures for metabolite ions detected through high-throughput exometabolomics remains limited. To obtain a clearer picture of the metabolites constituting individual and community substrate utilisation profiles we undertook an orthogonal methodology to assess metabolic niche occupancy of the SynCom strains, by measuring their substrate utilisation profiles using BIOLOG EcoPlate. We first determined the substrate utilisation profiles of all seven individual strains and the C7 (Table S6). The heatmaps derived from *in vitro* and *in silico* clearly discriminated the strains according to distinct metabolic niches (Fig 4a, b). *In vitro*, Elu prefered carbohydrates, Cin polymers and Ppu amines and amides. *In silico*, Elu also occupied a carbohydrate niche, while Ppu and Elu shared the amines and amide niche and Elu overall displayed the broadest capabilities, growing on 80% of the provided carbon sources (Fig 4b). Both *in vitro* and *in silico,* the C7 combined the substrate utilisation capacities of all constituent strains, illustrating the ‘addition effect’ that was consistently observed throughout this study. As observed before, Bpi did not grow on the defined medium *in silico,* which might be attributed to a difference in the BIOLOG EcoPlate medium (Table S7) and the *in silico* M9-based medium (Table S2).

**Figure 4:**
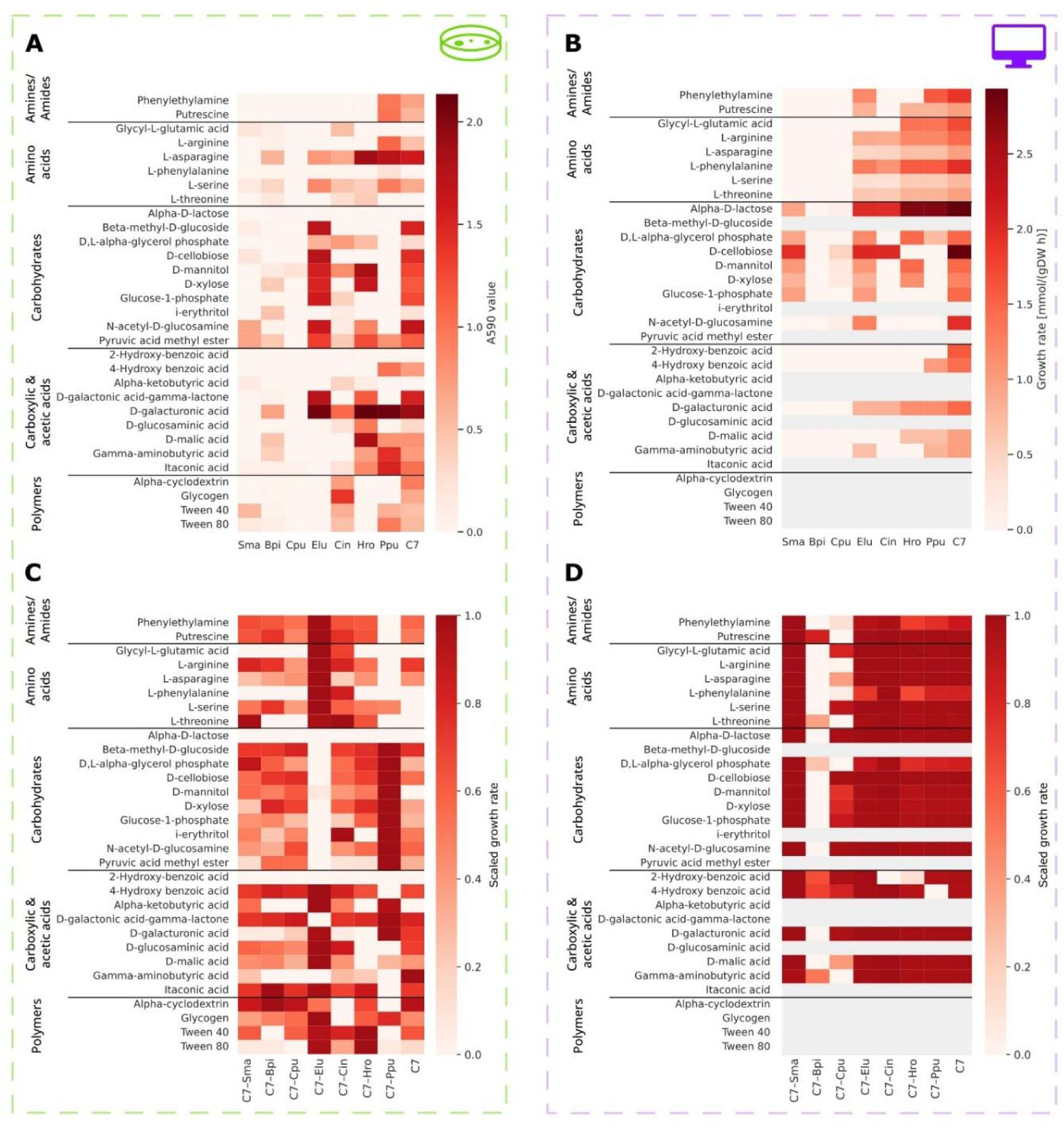
Metabolic niche profiling of individual strains, the C7 and C6 communities via phenotype microarray. (A) Heat map of substrate utilisation on 31 carbon sources by the SynCom strains measured *in vitro* using BIOLOG EcoPlates. Cell colour represents mean A590 value of three independent replicates. 2-Hydroxy-benzoic acid and Alpha-D-lactose were not utilised by any bacterial inoculation and excluded from the analysis. (B) Heat map of *in silico* substrate utilisation using the 31 carbon sources from the BIOLOG EcoPlate together with an M9 medium. Cell colour represents the predicted growth value with pFBA. A grey cell colour indicates the absence of a metabolite in the model due to missing exchange reactions. Cell colour represents predicted growth rates rather than substrate utilisation, as uptake rates were often upper-bound constrained and thus less informative. The mean growth rate of all seven bacteria in the community is used for the C7 scenario. (C) Heat map of substrate utilisation for the C6 communities (single strain removals) and the intact C7 SynCom measured *in vitro* using the BIOLOG EcoPlate. Data were background-corrected (blank well) and normalised to average well colour development (AWCD), followed by row-scaling across communities. Cell colour represents mean values of three independent replicates. (D) Heat map of *in silico* growth on the BIOLOG EcoPlate carbon sources in combination with an M9 medium. Values were row-scaled across communities. Cell colour represents predicted growth rates. For each community, the mean growth rate across all six or seven members was used. Original EcoPlate metabolite names are shown for *in silico* data in (B) and (D) while corresponding BiGG identifiers are listed in Table S7.

Then we assessed how much influence each strain exerts over the metabolic activity of the C7. We undertook C7 drop-out experiments analogous to the approach of Niu et al. [11], by removing individual strains from the community. *In vitro* results showed that removal of strains Elu and Ppu made the biggest impact on substrate utilisation. Specifically, removal of Elu manifested in lower utilisation of carbohydrates, whereas Ppu removal resulted in lower utilisation of amino acids and nitrogenous compounds (Fig 4c). *In silico* drop-out experiments revealed limited differentiation between most communities. Predicted growth rates were largely similar across substrates and chemical classes, which suggests a high degree of functional redundancy in the predicted substrate utilization (Fig 4d). Two exceptions were observed for C7-Bpi and C7-Cpu, which deviated from the remaining communities. When we further explored strain-specific effects by performing PCA on the predicted uptake and secretion fluxes of community simulations on the *in silico* maize root medium, C7-Elu showed the strongest deviation from all other drop-out communities (Supplementary Figure S2). These patterns mirror our *in vitro* observations and support a central metabolic role for *E. ludwigii*, consistent with its proposed keystone function in the SynCom.

### Cross-feeding phenotypes of the SynComs strains

We next explored pairwise cross-feeding interactions amongst SynCom strains. This initially involved determining which of the seven strains could grow on a minimal medium without any additional vitamins (Fig 5a, Table S8). Across nine carbon substrates, only Elu, Hro and Ppu significantly grew *in vitro*. This was also the case for the *in silico* simulations, except that Cin also showed a non-zero growth rate for most carbon sources (Fig 5b). Elu exhibited the highest growth rates *in vitro*, while *in silico* it showed growth across the widest range of carbon sources but only reached the highest growth rate for a single substrate (Fig. 5a, b and Supplementary Fig. S3).

**Figure 5:**
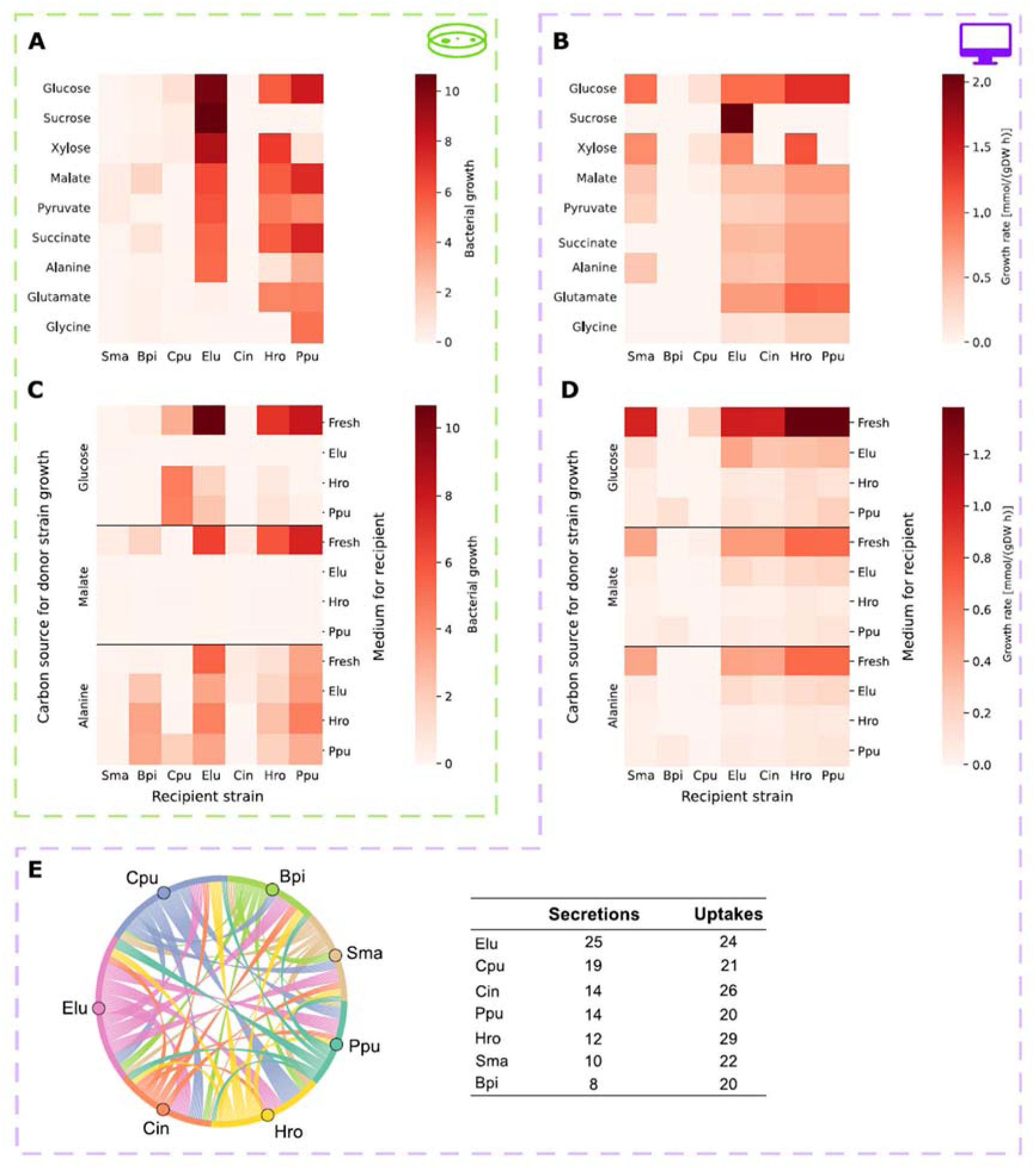
Growth assays and cross-feeding phenotypes of SynCom strains on minimal media. (A) Heat map of growth phenotypes for the individual SynCom strains cultivated on minimal media using nine sole carbon sources. Colour intensity corresponds to mean growth performance, measured via area under the curve in three independent experiments. (B) Corresponding *in silico* growth predictions under the same conditions as (A), with colour intensity indicating predicted growth rates. (C) Heat map of cross-feeding phenotypes for the seven SynCom strains grown on culture supernatants harvested from Elu, Hro and Ppu strains, following their pre-cultivation on either glucose, malate or alanine as sole carbon source. (D) Corresponding *in silico* cross-feeding based on predicted secretions from Elu, Hro and Ppu simulated under the same carbon source conditions as in (C), with colour intensity indicating predicted growth rates. (E) Chord plot illustrating qualitative metabolite exchanges within the C7, indicating which bacterium secretes a compound that is subsequently taken up by another community member. Connecting lines are coloured according to the secreting strain. A metabolite may therefore appear multiple times if utilised by several members. The accompanying table summarises the number of unique secretion and uptake reactions. Only exchanges with a flux < 0.01 mmol/(gDW h) were analysed.

Overall, Elu, Hro and Ppu were defined as prototrophic and our next step was to conduct cross-feeding assays, by harvesting filter-sterilised culture supernatants from the three prototrophic strains to provision as growth media for all seven strains (Fig 5c, Table S9). This dataset revealed that the effectiveness of cross-feeding involves a complex interplay of effects relating to donor strain, recipient strain and carbon source. For instance, the organic acid malate did not support cross-feeding *in vitro* in any of the studied strains, whereas the amino acid alanine was relatively effective at supporting cross-feeding for five recipient strains. There is little evidence of ‘donor effect’, because spent media harvested from all three prototrophic strains had similar potential for nourishing the recipients. When analysing the ‘recipient effects’, the ‘growth boost’ received by Bpi was particularly noteworthy. Bpi could not grow on fresh alanine medium, but grew effectively on culture supernatant harvested from three other SynCom members (Fig 5c). Complementary, we used computationally predicted secretion profiles as an *in silico* medium (see Methods). The results qualitatively reflected the *in vitro* patterns, with the difference that growth of the recipients was predicted to be more dependent on the donor strain itself than the carbon source for donor strain growth (Fig 5d). The compounds released by Elu enabled highest growth rates of recipient strains across all cross-feeding scenarios. Noteworthy, *in silico* Bpi grew on Ppu spent media, which led us to further investigate the cross-feeding fluxes within the C7 community model. The chord plot in Figure 5e shows that certain bacteria contribute more to the community through a high number of secretions, whereas others exhibited a broader range of uptakes. Elu secreted the largest number of metabolites (25) of which approximately 40% were amino acids. Among the 14 metabolites excreted by Ppu was thiamine (vitamin B1), which is taken up Bpi and thus enabled growth in this specific medium.

Taken together, these data provide experimental and computational evidence that cross-feeding occurs in this SynCom, and categorises the strains into three donors (Elu, Hro and Ppu) and two key recipients: Bpi and Cpu, the latter, interestingly, only *in vitro* analyses.

### B-vitamin dependencies of the SynCom strains

Our *in silico* results suggested thiamine dependency in Bpi. Literature evidence shows that B-vitamin auxotrophy is widespread among bacteria, positioning B-vitamin exchange as a critical mechanism for maintaining diversity within microbial communities [9]. Therefore, we investigated the B-vitamin responses of all seven SynCom strains, by studying their growth either with or without the addition of an eight-vitamin mixture containing all B-vitamins, in media containing a diversity of simple carbon substrates (Fig 6a, Table S10). *In vitro* data indicate that the strains can be grouped into three general categories: 1) High-responders (Bpi and Cpu) that receive a growth boost from B-vitamins; 2) Non-responders (Sma, Elu, Hro and Ppu) that grow strongly regardless of vitamin addition, and 3) Unknown (Cin) that did not grow in either condition. To quantify the magnitude of these differential vitamin responses, we measured growth curves for two exemplary strains (Bpi and Elu). Elu grows comparably under the two conditions, whereas Bpi receives a dramatic growth benefit from exogenous vitamins (Suppl. Fig. S4a).

**Figure 6:**
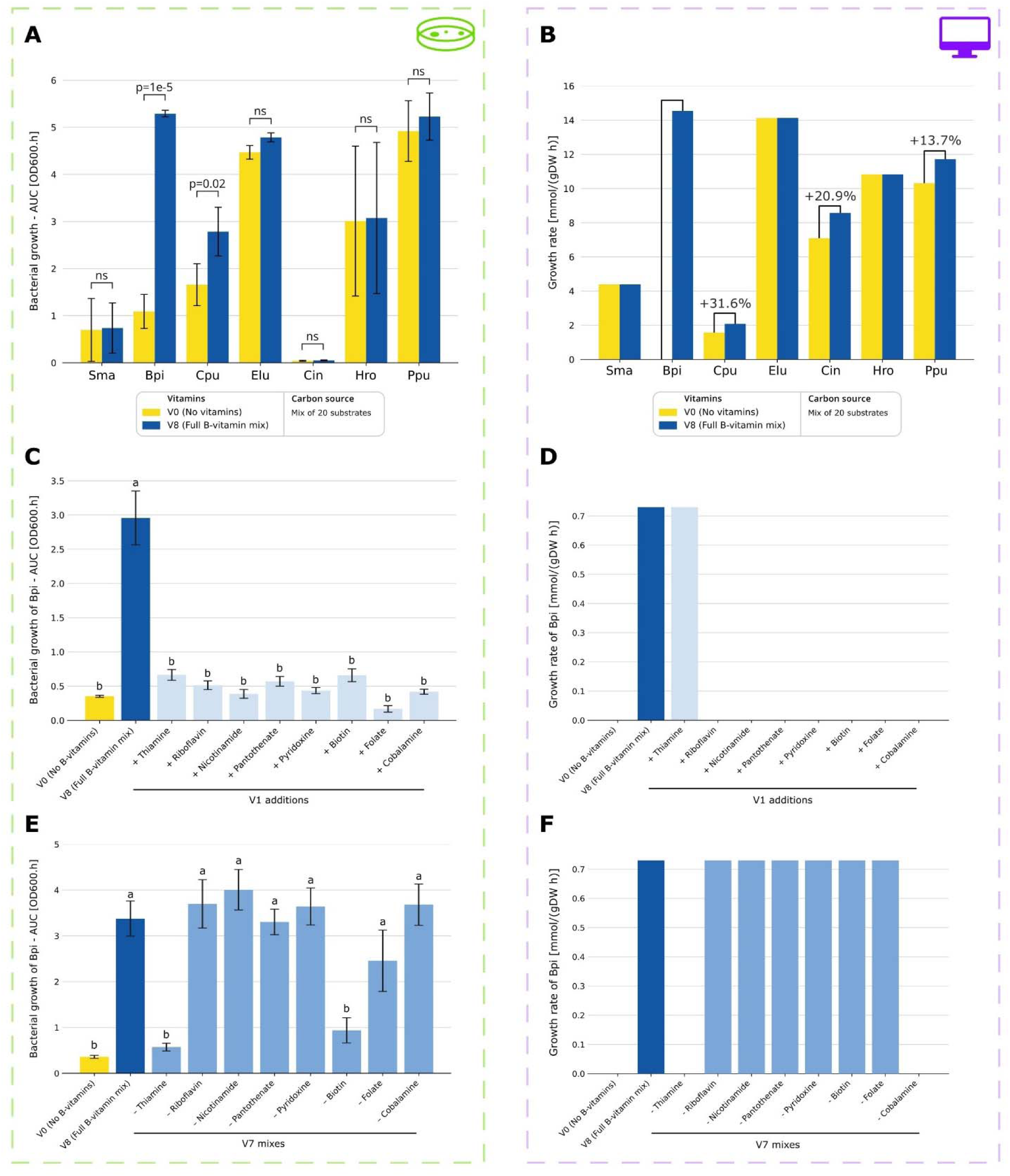
B-vitamin responses of the SynCom strains and pinpointing B-vitamin requirements for Bpi. (A) Bar chart showing the growth performance of each strain with or without B-vitamin mixture. Statistical significance of each strains’ vitamin response was determined using Student’s t-test. Error bars represent SD, n=4. (B) Bar chart showing the predicted growth rates of each strain with or without B-vitamin mixture. (C) Bar chart showing the effect of adding single B-vitamins on Bpi growth performance. Groups annotated with the same letter are not significantly different using Tukey’s HSD test (α=0.95). Error bars represent SD, n=4. (D) Bar chart showing the effect of adding single B-vitamins on simulated growth rates of Bpi. (E) Bar chart showing how Bpi’s growth performance is affected by V7 dropout mixes. Groups annotated with the same letter are not significantly different using Tukey’s HSD test (α=0.95). Error bars represent SD, n=4. (F) Bar chart showing how Bpi’s simulated growth rate is affected by V7 dropout mixes.

Overall, this profile of vitamin dependency (Fig. 6a) is largely consistent with the results of our cross feeding assays (Fig 5). Across the two datasets, the same strains which were grouped as ‘cross-feeding donors’ (Elu, Hro and Ppu) showed no requirement for exogenous vitamins, whereas the strains that were grouped as ‘key recipients’ (Bpi and Cpu) received a growth benefit from the addition of a vitamin mixture. The *in silico* picture looks similar; Sma, Elu and Hro are non-responders of which Elu and Hro have much higher growth rates than Sma (Fig 6b). Bpi, Cpu, Cin and Ppu respond to the vitamin addition and in case of Bpi this enables growth. The latter support the notion of Bpi as recipient strain within the community.

Taken together, experimental evidence and metabolic modelling suggest that vitamin exchange between prototrophs and auxotrophs belongs to mechanisms mediating stability of this SynCom.

We then undertook reductionist experiments to pinpoint which vitamins were required by the auxotrophic strain Bpi. We first hypothesised, based on computational predictions, that Bpi was auxotrophic for one specific B-vitamin, but the provision of any single vitamin did not rescue Bpi’s growth phenotype *in vitro* (Fig 6c). Our next step was to undertake V8 ‘drop-out’ experiments, by removing one single vitamin from the 8-vitamin mixture. Results of this experiment revealed that a significant drop in Bpi’s growth performance was elicited by two of the V7 mixes (V8-Thiamine and V8-Biotin) (Fig 6e). This strongly suggested that Bpi is a double auxotroph for thiamine and biotin. We corroborated this finding by showing that addition of these two vitamins rescued Bpi’s growth phenotype in a manner like the V8 (Supplementary Fig. S4b). Interestingly, computational V1 and V7 simulations for Bpi could only reproduce the auxotrophy for thiamine and not biotine (Fig. 6 d and f). Taken together, we conclude that Bpi is auxotrophic for both thiamine and biotin, which therefore positions the exchange of these metabolites as one candidate mechanism mediating the stability of this SynCom.

## Discussion

The motivation of this study was to define specific metabolic mechanisms that mediate strain coexistence in an established bacterial SynCom, by exploring the processes of niche differentiation and cross-feeding. Using multiple methods, we comprehensively documented the metabolic niches occupied by each strain, which demonstrated that the SynCom members are metabolically complementary. Furthermore, we described which specific strains are the donors and recipients of cross-feeding interactions, and showed that some strains were auxotrophic whereas others were prototrophic. Results of vitamin dependency experiments implicated certain B-vitamins as candidate molecules exchanged between strains. Interpreting our results, we are particularly interested in defining metabolic traits which could serve as ‘design principles’ for rationally assembling stable SynComs.

### Metabolic complementarity as a mechanism underpinning SynCom stability

The ecological principle of niche differentiation states that species which coexist in the same habitat must consume different resources in order to avoid competition [30]. This principle could guide efforts to compose stable microbial communities for applications in agriculture and medicine, by matching compatible strains according to their non-overlapping substrate preferences [1]. The SynCom investigated in this study was previously shown to assemble stably on maize roots [11], and therefore, we postulated that the constituent strains would occupy complementary metabolic niches.

Using multiple experimental and computational approaches, we conclusively demonstrate that the strains in this SynCom exhibit metabolic niche complementarity. This is shown by three orthogonal lines of evidence: 1) Metabolomic footprinting on maize root extracts, which pinpointed hundreds of metabolites only consumed by individual strains, tentatively matched to maize secondary metabolites; 2) Phenotyping microarrays revealing that the strains utilised different classes of metabolic substrates (ie: polymers, carbohydrates, amides); and 3) Vitamin auxotrophy profiling, which clearly distinguished between a set of three prototrophic strains versus two auxotrophs. Consistent with the experimental data, *in silico* analyses independently identified complementary metabolic strategies across strains, indicating that niche differentiation is encoded in their metabolic networks. Interpreting these findings, we postulate that metabolic complementarity could be a causative mechanism mediating stability in this SynCom. Potentially, future experiments could test this principle across a larger number of differentially formulated SynComs. For example, would SynComs composed of strains with overlapping substrate preferences exhibit a collapse in diversity due to competitive exclusion effects? In turn, can SynCom stability be promoted by combining strains that occupy non-overlapping metabolic niches? And would it be possible to engineer interdependence between strains, by combining vitamin exporters with reciprocal auxotrophs?

In the literature, the primary rationale guiding SynCom assembly has been taxonomic representativeness, whereby individual SynCom strains are selected as representative members corresponding to the major phylogenetic groupings which occur in naturally assembled microbiome [31]. It is often presumed that taxonomically diverse communities will inevitably exhibit functional diversity, because the different microbial phyla have committed to diverging ecological strategies early in their evolutionary history [32,33]. However, there is increasing evidence that bacterial metabolic traits are poorly predicted by taxonomy, with closely related strains often showing widely diverging substrate preferences, potentially mediated the horizontal transfer of metabolic genes [34]. In our opinion, the concept of metabolic complementarity could be a useful tool for designing stable SynComs, because it has a stronger mechanistic basis for promoting stability compared to the standard approach of combining diverse taxa. *In silico* screening of candidate communities via metabolic modelling offers a time and cost-efficient means to prioritise metabolically complementary strain combinations for subsequent experimental testing *in vitro*.

### New insights into the metabolic properties of a keystone strain

Keystone strains are of major interest in the field of microbiome engineering, because the presence of these strains promotes the diversity of the surrounding community. Therefore, keystone strains could potentially be administered in cases of microbiome dysbiosis to restore community diversity or alternatively included into SynComs to promote the coexistence of accompanying strains [35]. There is significant interest in defining which members of the plant microbiota are keystones [36], and the work of Niu et al [11] provided a useful resource to the field, because it empirically defined Elu as a keystone strain stabilising SynCom assembly on maize roots. Here, our results build upon this previous work, by providing new mechanistic information about the metabolic properties of Elu in relation to the other SynCom strains and the C7.

In this study we show that Elu is a fast-growing strain with a broad metabolic niche and a large metabolic influence on the C7. It can rapidly utilise a diverse range of carbon substrates without the need for exogenous vitamins, and its culture supernatants can sustain other strains in cross-feeding assays. We postulate that these results provide a mechanistic explanation for the keystone behaviour of Elu demonstrated in Niu et al [11]. Specifically, one element of Elu’s keystone behaviour could be its overlapping metabolic niche with the disruptive Cpu (Fig 2). Potentially, the faster-growing Elu could outcompete Cpu for the primary usage of these maize secondary metabolites, which could prevent Cpu from dominating the community because these plant-substrates could serve as building blocks for antimicrobial compounds synthesised by Cpu. Our *in silico* studies identified 3-hydroxydecanoic acid, which has pathogen suppression and plant immunity activation properties, as one potential metabolite [29].

A further element could be Elu’s cross-feeding ability, whereby this prototrophic strain synthesises vitamins that are subsequently released into the environment to promote the coexistence of auxotrophic strains. Future experiments could test whether the metabolic properties exhibited by Elu are general characteristics shared by all keystone strains, or whether they are only relevant in the context of this simplified SynCom. If these metabolic characteristics are indeed common amongst keystones, then potentially they could be used as criteria for identifying candidate keystone strains from large panels of microbial isolates.

### Tailoring stable SynComs for specific plant genotypes

Plant species differ in their root chemistry [37], and there is growing evidence that species-specific secondary metabolites act as a selection of pressure shaping the composition of the rhizosphere microbial community [38]. This has an impact on the design of plant-associated SynComs, because microbial strains need to be equipped to colonise the chemical environment unique to the target host plant. The C7 SynCom [11] was assembled via host-mediated selection on maize roots, which automatically means that these strains are competent in the maize root environment. In this study, we provide new information about the metabolic niches occupied by each strain, using LC-MS exometabolomics to infer that maize secondary metabolites represent a major source of differential metabolic niches, particularly for the strains Elu and Cpu. However, we have not authenticated the identity of these maize root metabolites using reference standards, so further investigations are necessary before we can unequivocally identify the specific metabolites that nourish each strain. Nevertheless, the exometabolomic approach utilised in this study can be a useful tool in tailoring SynCom design for distinct plant species, because it can facilitate the optimal matching between host root chemistry versus microbial substrate utilisation. In parallel, *in silico* metabolic modelling enables the systematic identification of candidate metabolites predicted to be secreted or exchanged between community members, provided that a representative exudate profile for the host species is available. Moreover, once such exudate compositions are defined, modelling allows rapid exploration of alternative plant species or genotypes, thereby supporting the rational design and prioritisation of plant specific SynComs for experimental validation.

Metabolic modelling provides a complementary framework to interpret the experimental observations, while remaining a simplified abstraction of the maize root environment. The simulations were based exclusively on primary metabolites identified by GC-MS and therefore do not capture the full complexity of maize root exudates, particularly secondary metabolites that likely contribute to niche differentiation in planta. Model predictions are consequently most informative when media compositions are well defined, and incorporating more comprehensive, condition-specific exudate profiles will enhance the biological relevance of future simulations. Furthermore, flux balance analysis relies solely on genome-encoded metabolic potential and does not account for regulatory effects, enzyme kinetics, or gene expression, which may explain discrepancies observed for some substrate utilisation and community-level phenotypes. Despite these limitations, the approach enables rapid, systematic hypothesis testing and low-cost exploration of complex metabolic conditions. While model-guided experimental testing is well established for single organisms, experimentally validated community models remain rare. Here, community-level model predictions have been directly confirmed through experiment, highlighting the broader potential of this approach for microbiome research. The qualitative agreement between predicted and observed metabolic niches, vitamin dependencies, and the central role of *E. ludwigii* supports the utility of genome-scale community modelling as a tool to contextualise and guide experimental analyses.

### Conclusion

The characterisation of metabolic phenotypes of each member of a seven-strain bacterial SynCom, established by Niu et al. [11], defines the metabolic interplay between the strains by documenting their differential metabolic niche occupancies and exploring donor-recipient cross-feeding interactions. We postulate that these metabolic mechanisms could be illustrative of general principles underpinning the stability of microbial communities, which could be used to guide the rational assembly of stable SynComs.

## Declarations

### Ethics approval and consent to participate

Not applicable

### Consent for publication

Not applicable

### Availability of data and material

All data generated or analysed during this study are included in this published article and supplementary material. The code needed to reproduce the results of this paper can be found at https://github.com/Toepfer-Lab/C7-maize-syncom.

### Authors’ contributions

R.P.J. initiated the study and supervised the experimental part. R.P.J., S.K., and N.T. designed the experiments. J.K. performed the experiments, assisted by P.K. in the analyses of carbon sources. V.W., S.M., and T.I. performed metabolomics analyses. J.K. and R.P.J. analyzed the experimental data. L.S. performed the modelling supported by M.F., and supervised by N.T.. J.K., L.S., N.T., S.K. and R.P.J. prepared figures and wrote the manuscript. All authors reviewed and approved the final version.

## Acknowledgements

We thank Ben Niu (Northeast Forestry University, China) for providing bacterial strains, Nicole Mantke (Institute of Geology and Mineralogy, University of Cologne) for performing TOC measurements, and the Service Unit for Metabolomics and Lipidomics at the University of Göttingen for GC-MS access.

## Supplementary Information

### Supplementary Figures

**Supplementary Figure S1:** Enrichment of metabolites by the SynCom and constituent strains.

**Supplementary Figure S2:** Metabolic analysis of C6 drop-out communities via PCA.

**Supplementary Figure S3:** Growth assays of SynCom strains on minimal media.

**Supplementary Figure S4:** B-vitamin responses of the Bpi and Elu strains.

### Supplementary Tables

**Table S1:** Abundance values for 25 primary metabolites present in maize root extract and depleted by bacterial inocula in GC-MS exometabolomic experiments.

**Table S2:** Composition of M9 medium.

**Table S3:** Abundance values and LC-MS information for 425 metabolite ions present in maize root extract and depleted by at least one bacterial inoculum in LC-MS exometabolomic experiments.

**Table S4:** Abundance values and LC-MS information for 228 metabolite ions enriched in the extracellular medium by at least one bacterial inoculum in LC-MS exometabolomic experiments.

**Table S5:** Composition of the *in silico* maize root medium.

**Table S6:** Substrate utilisation measurements for 29 carbon sources present on BIOLOG Ecoplate phenotype microarrays.

**Table S7:** *In silico* metabolites used to represent the Biolog Ecoplate carbon sources.

**Table S8:** Growth performance of the individual SynCom strains measured using *in vitro* growth assays on nine different metabolic substrates provided as single carbon source in M9 medium.

**Table S9:** Growth performance of the individual SynCom strains measured using *in vitro* cross-feeding assays, where two factors were systematically manipulated: the carbon source and the medium history.

**Table S10:** Growth performance of the individual SynCom strains measured using *in vitro* assays to determine overall vitamin auxotrophy.

**Table S11:** Growth performance of strain Bpi measured using *in vitro* assays to determine which specific vitamins are required for growth.

