## Supplemental Methods for "Metabolic profiling and genome-scale modelling uncover mechanistic drivers of microbiome stability in synthetic maize root community"

**Supplementary Information**

**Reconstructing and curating metabolic models of bacteria**

Genome scale metabolic models were reconstructed using CarveMe (Machado et al. 2018), generating draft models from protein FASTA files. Reconstructions were performed using Gram specific parameters and soft constraints to exclude reactions listed in a TSV file (https://github.com/Toepfer-Lab/C7-maize-syncom/blob/main/Models/01_carveme/exclude_reaction.tsv). This exclusion list was derived from the iYO844 model (Oh et al. 2007), as initial unconstrained reconstructions introduced duplicate and biologically implausible reactions. Consequently, reactions unique to this model and not supported by other BiGG database entries were removed. CarveMe outputs were generated in SBML (XML) format and analysed using COBRApy.

Model quality was assessed using MEMOTE (Lieven et al. 2020), and reports guided subsequent curation. Mass and charge imbalances were resolved through a two-step procedure. First, metabolite annotations were standardised by overwriting model entries with BiGG database information. Second, remaining imbalanced reactions were manually curated by comparison with cross referenced databases including KEGG and MetaCyc, adjusting formulas and charge states accordingly.

Duplicate reactions and metabolites were further analysed using MACAW (Moyer et al. 2025), which classifies duplicates into exact, directional, coefficient, and redox categories. Exact and directional duplicates were resolved automatically using a custom Python script that integrates gene protein reaction rules to determine appropriate merging. Remaining cases were manually inspected, while coefficient and redox variants were retained where biologically justified.

The universal biomass reaction generated by CarveMe, which is not organism specific, was evaluated by testing the producibility of each biomass component via flux balance analysis. Model capabilities were compared with pathway information from KEGG PATHWAY and MetaCyc to determine whether components required removal, replacement, or pathway completion.

*In vitro* results of this study lead to the investigation of sucrose pathways. The comparison to computational results revealed transport reactions for sucrose originating from unreliable BiGG models causing excessive growth rates. Consequently, reactions with the IDs SUCpts, SUCRt2 and EX_sucr_e were removed from the models. To prevent the creation of orphan metabolites additional reactions and metabolites were subsequently eliminated.

To approximate the maize root environment, a simulation medium was defined based on metabolites identified by GC-MS analysis (see Suppl. Table S3, S5). Not all compounds could be matched to corresponding BiGG IDs, and some metabolites are absent from the models. On the contrary, some metabolites have multiple equivalents in BiGG due to isomers. In addition to a maize root extract, simulations included an additional M9 minimal medium to provide salts, ions and a nitrogen source (see Suppl. Table S6). The composition is based on the M9 medium used in the *in vitro* analyses of this study. All M9 components, were assigned a high uptake bound of 1000.0 mmol/(gDW h) to prevent artificial growth limitations. In contrast, carbon sources from the maize root extract were constrained to 10.0 mmol/(gDW h). This value is widely applied in genome-scale metabolic modelling as a physiologically realistic upper limit for substrate uptake, reflecting the fact that carbon uptake in bacteria is inherently rate-limited by transport and enzymatic capacities. This approach is consistent with established practices in *E. coli* and other microbial metabolic reconstructions (Feist, Monk, Machado, Corrao).

**Tentative Metabolite Annotation of MS-Features**

To assign candidate IDs to the metabolite ions of interest, m/z values were searched using CEU Mass Mediator (Gil-de-la-Fuente et al., 2019) against the Metlin database (Smith et al., 2005) with 10 ppm tolerance, for various common adducts (positive mode: M+H, M+Na, M+K; negative mode: M-H, M+Cl, M+FA). In cases where multiple database compounds matched to the same m/z value, the compound with the lowest Metlin database identifier was chosen. These candidate metabolite IDs were then grouped into chemical categories using the ClassyFire database (Feunang et al., 2016).
