## Supplemental Figures for "Metabolic profiling and genome-scale modelling uncover mechanistic drivers of microbiome stability in synthetic maize root community"

**Supplementary Figures**

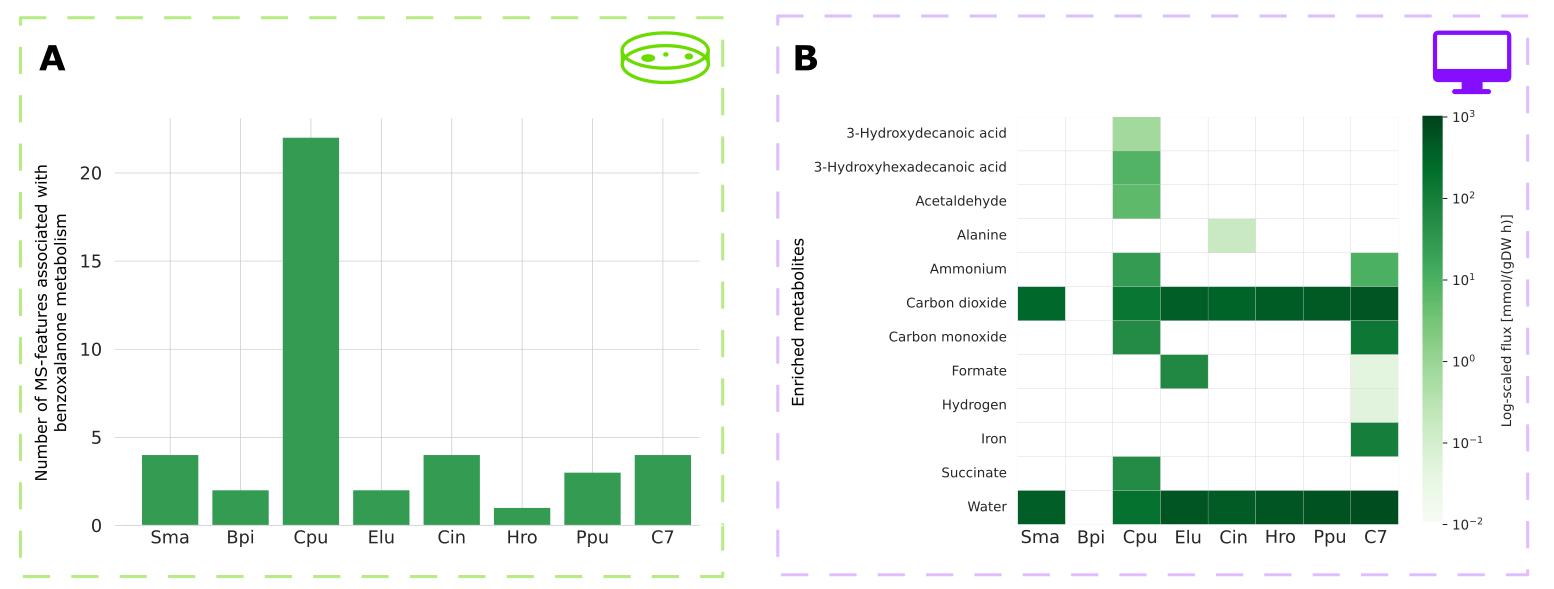
***Supplementary Figure S1: Enrichment of metabolites by the SynCom and constituent strains.***

(A) Bar chart showing count of LC-MS features tentatively matched to benzoxalanone compounds across strains. Features were assigned based on matches to the ClassyFire category organoheterocyclic compounds. Annotation is at the MS1 level only without use of confirmation standards. Full measurement details are given in Supplementary Table S3. (B) Heat map of all secreted metabolites by the SynCom strains and the community with a flux > 0.01 mmol/(gDW h). All models were simulated on a medium consisting of the primary metabolites identified via GC-MS and a M9 medium. Cell colour represents the flux value of the exchange reaction transporting a metabolite into the extracellular compartment***.***

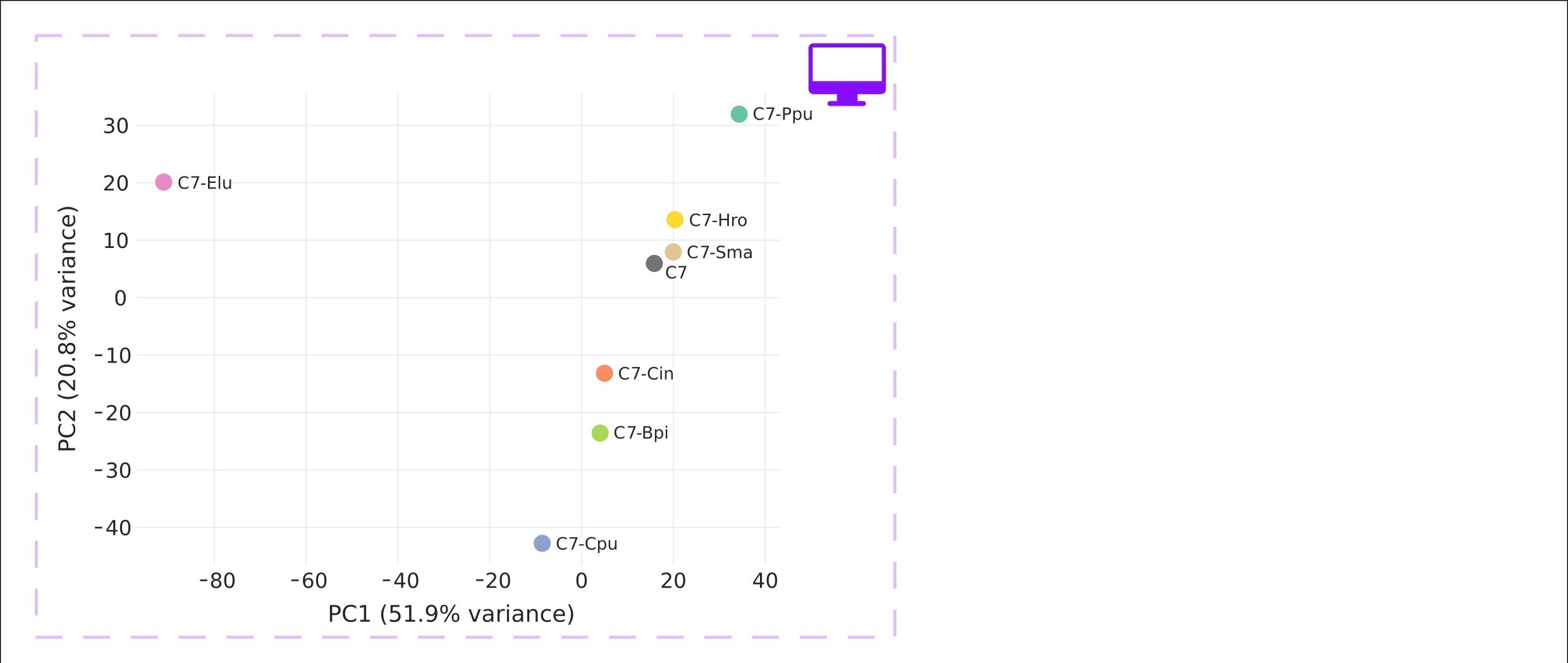

***Supplementary Figure S2: Metabolic analysis of C6 drop-out communities via PCA.***

PCA of all fluxes from exchange reactions predicted with pFBA. All models were simulated on a medium consisting of the primary metabolites identified via GC-MS and a M9 medium. Community names indicate which member was dropped to create a six-member community.

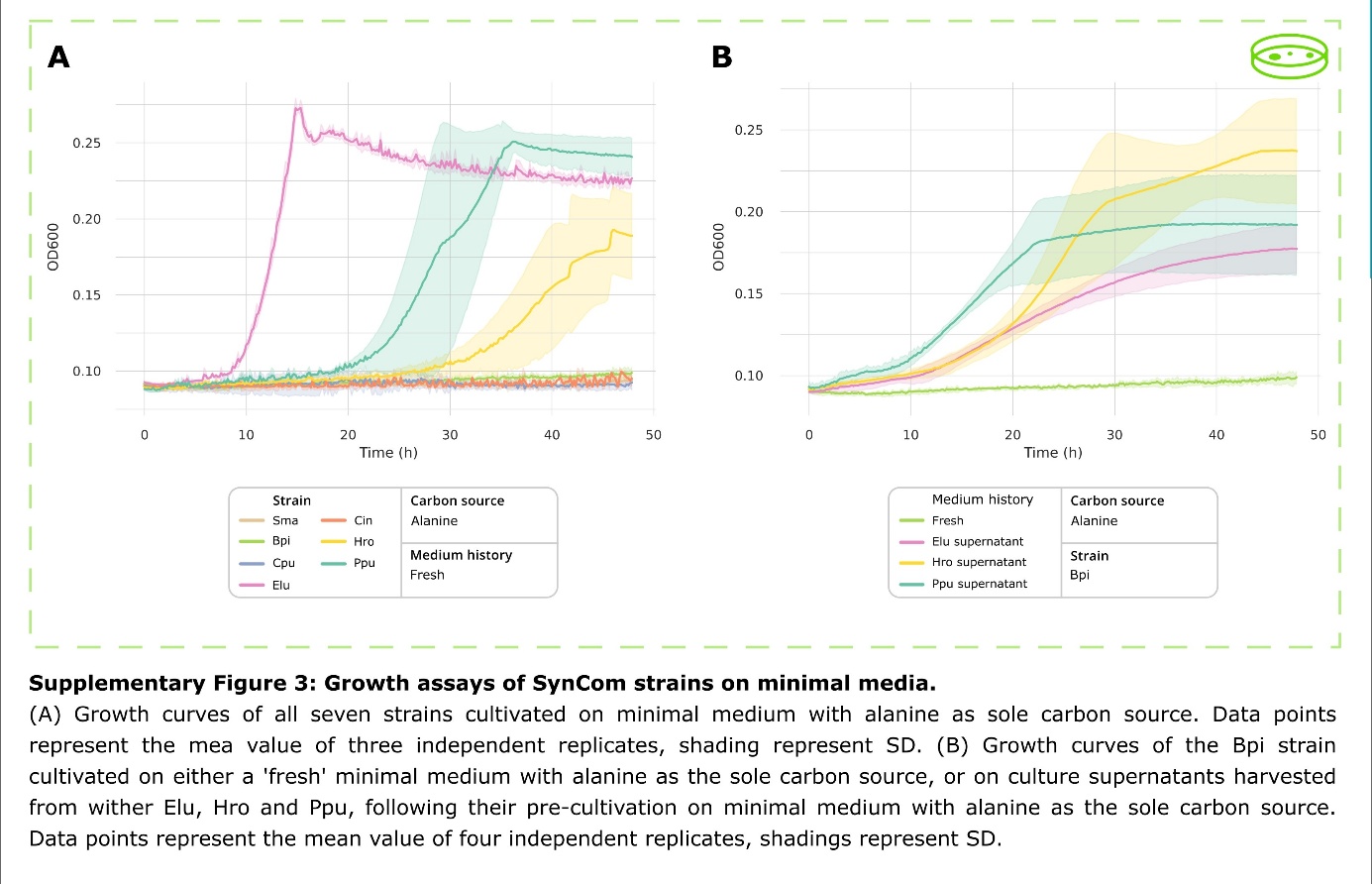

***Supplementary Figure S3: Growth assays of SynCom strains on minimal media.***

(A) Growth curves of all seven strains cultivated on minimal medium with alanine as sole carbon source. Data points represent the mean value of three independent replicates, shading represent SD. (B) Growth curves of the Bpi strain cultivated on either a 'fresh' minimal medium with alanine as the sole carbon source, or on culture supernatants harvested from wither Elu, Hro and Ppu, following their pre-cultivation on minimal medium with alanine as the sole carbon source. Data points represent the mean value of four independent replicates, shadings represent SD.

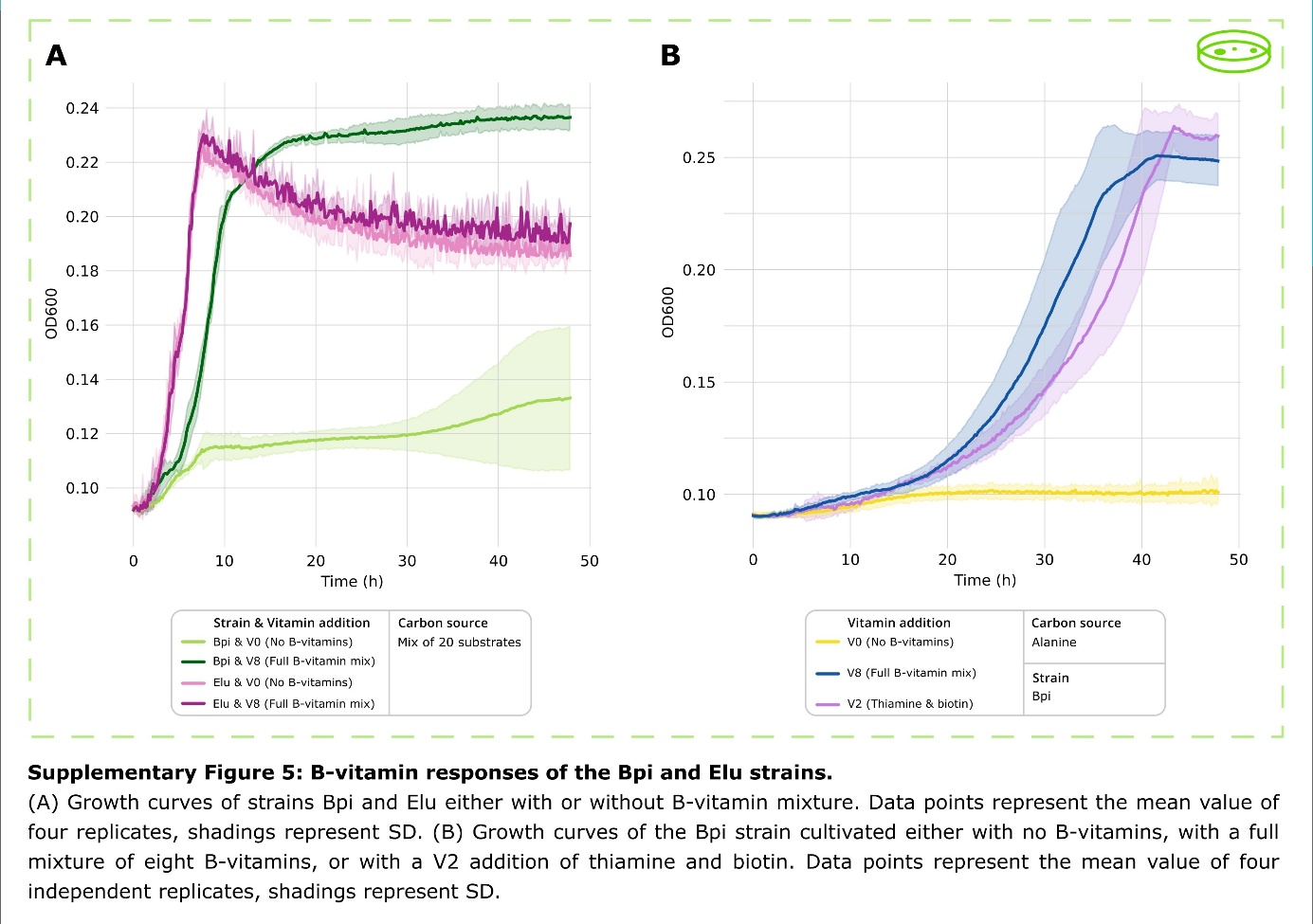

***Supplementary Figure S4: B-vitamin responses of the Bpi and Elu strains.***

(A) Growth curves of strains Bpi and Elu either with or without B-vitamin mixture. Data points represent the mean value of four replicates, shadings represent SD. (B) Growth curves of the Bpi strain cultivated either with no B-vitamins, with a full mixture of eight B-vitamins, or with a V2 addition of thiamine and biotin. Data points represent the mean value of four independent replicates, shadings represent SD.
